# Contextual experience modifies functional connectome indices of topological strength and organization

**DOI:** 10.1101/2020.06.13.150060

**Authors:** Marjory Pompilus, Luis M. Colon-Perez, Matteo M. Grudny, Marcelo Febo

**Author notes:** **Corresponding author:** Marcelo Febo, PhD, P.O. Box 100256, Department of Psychiatry, College of Medicine, University of Florida, Gainesville, FL 32611.

## Abstract

Stimuli presented at short temporal delays before functional magnetic resonance imaging (fMRI) can have a robust impact on the organization of synchronous activity in resting state networks. This presents an opportunity to investigate how sensory, affective and cognitive stimuli alter functional connectivity in rodent models. In the present study, we assessed the effect of a familiar contextual stimulus presented 10 minutes prior to sedation for imaging on functional connectivity. A subset of animals were co-presented with an unfamiliar social stimulus in the same environment to further investigate the effect of familiarity on network topology. Female and male rats were imaged at 11.1 Tesla and graph theory analysis was applied to matrices generated from seed-based functional connectivity data sets with 144 brain regions (nodes) and 10,152 pairwise correlations (edges). Our results show an unconventional network topology in response to the familiar (context) but not the unfamiliar (social) stimulus. The familiar stimulus strongly reduced network strength, global efficiency, and altered the location of the highest eigenvector centrality nodes from cortex to the hypothalamus. We did not observe changes in modular organization, nodal cartographic assignments, assortative mixing, rich club organization, and network resilience. The results suggest that experiential factors, perhaps involving associative or episodic memory, can exert a dramatic effect on functional network strength and efficiency when presented at a short temporal delay before imaging.

## INTRODUCTION

Over the past two decades, the combined use of functional magnetic resonance imaging (fMRI) with behavioral and cognitive paradigms has been a major driving force for research on the neural mechanisms of memory (1). These studies, largely in human subjects, have helped establish a role of neural processing in the hippocampal formation in learning while additional distributed cortical and subcortical neuronal populations are considered critical to long-term episodic memory storage and recall (2–4). Our view of the mechanisms of memory has further evolved through recent neuroimaging work applying graph theory to analyze brain wide functional connectivity data (5–8). Indeed, novel functional network properties have been uncovered in human subjects undergoing learning and memory tasks during scanning (9–12).

There has been much less work using network analyses in rodent models, particularly in relation to memory-related neural processing (13, 14). Many available studies that have investigated the neural mechanisms of learning and memory by applying resting state fMRI and functional connectivity analysis have focused on comparisons between young and aged male rat models. In adult male rats, functional connectivity between hippocampal CA3 subfield and thalamus, septum, sensoricortical and cingulate cortical regions is strengthened as a function of improved performance on the Morris water maze (MWM) (15). Performance on the MWM in aged male rats correlates with strength of default mode connectivity with the retrosplenial cortex (16), and weakening of CA1-prefrontal connectivity is associated age-related decline in learning (17). An index of rich club organization, which reflects the presence of a small group of very high degree ‘hub’ nodes, was observed to be greater in aged versus young male rats, and increased as a function of cognitive training (18). Longitudinal decline in memory was linked to changes in functional connectivity between the dorsal hippocampus and retrosplenial cortex, thalamus, striatum, and amygdala (19). In a study using optimal percolation theory to quantify the collective influence of hub nodes in male rat fMRI networks, Del Ferraro *et al* (2018) observed that hippocampal CA3-CA1 long-term potentiation (LTP) caused a potent global integration of brain BOLD activation which was weakened by elimination of a low degree node such as the nucleus accumbens (20). More neuroimaging research in animal models of learning and memory are needed to explore novel links between functional network topology and underlying synaptic and molecular mechanisms (21). Such studies are needed to confirm the utility of graph theory based quantifiers as biomarkers of brain function in control conditions, aging, and in neurological diseases.

A limitation to investigating functional connectomics of memory in rodent models is that behavioral paradigms for learning and memory can involve substantial training and variable temporal delays between testing and fMRI acquisition. As of yet, studies in rodents have not taken advantage of the fact that behavioral tasks (e.g., motor tasks) carried out minutes prior to resting state scanning can be remapped onto anatomically-relevant functional connectivity networks (22). Related to this experimental phenomenon, a previous study from our laboratory suggested that an unexpected arousing stimulus (i.e., handling during administration of a saline injection) given to rats 10 minutes prior to sedation and setup for fMRI acquisition significantly increased functional connectivity across multiple brain regions in rats (23). Thus, rats seemed to have retained a ‘memory’-like neural response of the transient pre-scan stimulus (23). In the present study we used a variation of this brief pre-scan stimulation approach to investigate the effect of a familiar stimulus (contextual environment) versus a memory-engaging unfamiliar stimulus (social interaction with a juvenile conspecific) on brain modularity, global and nodal strength, communication efficiency, and other quantifiers of network topology. Harnessing a full battery of connectomic calculations we observed that prior experience significantly suppresses functional network strength, reduced communication efficiency, and rearranged the location of putative central hub nodes from the cortex to subcortical areas. The results suggest that familiar stimuli engage cortical networks involved in memory more robustly than novel unfamiliar stimuli. Future studies will be needed to determine if the observed weakening of network strength and efficiency, and hub reorganization, under the present experimental paradigm may act to minimize interference by novel stimuli during reconsolidation processing of a familiar stimulus (24).

## MATERIALS AND METHODS

### Subjects

Thirty adult female and male Long Evans rats (225-350 g on arrival) were obtained from Charles River laboratories (Wilmington, NC). An additional 12 juvenile female and male rats (100-150 g upon arrival; 6 per sex) were used as social stimuli prior to imaging. Rats were housed in sex-matched pairs in a temperature and humidity-controlled vivarium with a 12 h light cycle (lights on at 0700 h) and food and water provided *ad libitum*. All procedures received prior approval from the Institutional Animal Care and Use Committee of the University of Florida and followed all applicable NIH guidelines.

**Figure 1.**
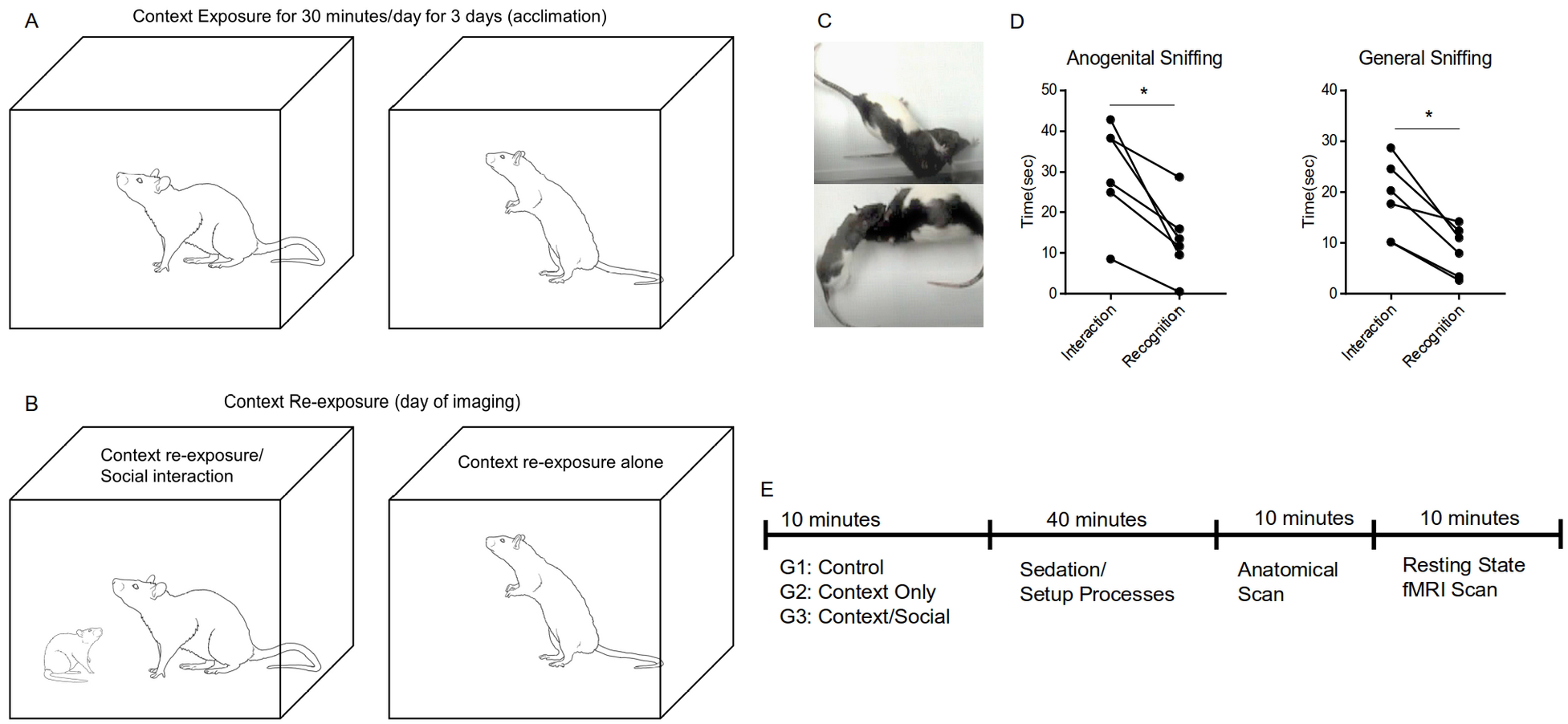
Timing of pre-scan stimulus presentation, sedation, setup, and image acquisition were kept the same across all subjects. A) Rats are placed into the same test cage over 3 days prior to imaging (become familiar). B) Before imaging, rats are once again presented with the familiar context either in the absence or presence of an unfamiliar social stimulus (juvenile rat). C) Digital video frames showing visually confirmed social interactions. D) Assessments of exploratory sniffing. Rats explore novel juvenile rats at a much higher level at the first 10 minute test than in a second test 1 hour later (paired t-test, p<0.05). E) Temporal sequence between stimulus presentation and fMRI acquisition.

### Experimental design

Rats were assigned to one of 3 experimental groups, which included a naïve *control* group (no pre-scan exposure; n=11, with 5 males), a *cage only* group (*context exposure* group; n=10, with 5 males), and a *cage+social stimulus* group (*context/social exposure*; n=9, with 5 males). The context and context/social exposure groups were acclimated for 30 minutes a day for 3 consecutive days prior to imaging to a clear plastic test cage (dimensions: 40 cm^3^) surrounded along the exterior of its walls with a black foam padding to minimize light inside the test cage (Figure 1A). Acclimation was carried under low noise and dim light conditions in the MRI facility room near the scanner. On the day of imaging a subset of context exposed rats were presented with an unfamiliar juvenile rat of the same sex inside the familiar test arena for 10 minutes (Figure 1B). Exploratory interactions between rats were visually confirmed (Figure 1C). In a cohort of rats tested separately but in the same test cages we observed that test rats interact with the unfamiliar rat through repeated bouts of exploratory sniffing, grooming, and pursuing inside the test cage (Figure 1D). Typically, during a second exposure 1 h later (which corresponds to the time in which rats are scanned) the test rats reduce exploratory behaviors towards the now familiar rat (Figure 1D). In the present study, each rat was scanned 50 minutes after initial social exposure (Figure 1E). For the context group, rats were exposed to an empty test arena (familiar context) for 10 minutes and imaged 50 minutes later (Figure 1E). To control for effects of context and context/social exposure, we included a control group of rats imaged without any experimental manipulation other than the same imaging setup process that rats in the other two groups underwent (Figure 1E). All handling of rats, acclimation, pre-scan exposures, and image data collection were carried out by the same investigator to minimize novelty stress effects.

### Magnetic resonance imaging

Images were collected in an 11.1 Tesla MRI scanner (Magnex Scientific Ltd., Oxford, UK) with a Resonance Research Inc. gradient set (RRI BFG-240/120-S6, maximum gradient strength of 1000 mT/m at 325 Amps and a 200 μs risetime; RRI, Billerica, MA) and controlled by a Bruker Paravision 6.01 console (Bruker BioSpin, Billerica, MA). A custom-made 2.5 cm × 3.5 cm quadrature radiofrequency (RF) surface transmit/receive coil tuned to 470.7MHz (^1^H resonance) was used for B1 excitation and signal detection (RF engineering lab, Advanced Magnetic Resonance Imaging and Spectroscopy Facility, Gainesville, FL). Rats were administered a subcutaneous injection of 0.025 mg/kg dexmedetomidine (Patterson veterinary) during setup and imaged under a continuous flow of 1% isoflurane (delivered at 0.1L/min mixed with medical grade air containing 70%N_2_ and 30% O_2_). Using this protocol, we previously reported that rats had stable breathing rates (45-65 beats per min), heart rate, and relative blood oxygen saturation for over 1 hour post-dexmedetomidine (25). Animals were placed on a custom plastic bed with a respiratory pad underneath the abdomen. Respiratory rates were monitored continuously, and body temperature was maintained at 37-38°C using a warm water recirculation system (SA Instruments, Inc., New York). For each rat we acquired a 10-minute high-resolution T2 weighted anatomical scan followed by a 10-minute functional magnetic resonance imaging (fMRI) scan without any stimulation (under ‘resting state’ conditions). Functional MRI scans were acquired at least 50 minutes after dexmedetomidine administration.

A T2-weighted Turbo Rapid Acquisition with Refocused Echoes (TurboRARE) sequence was acquired with the following parameters: effective echo time (TE) = 41 ms, repetition time (TR) = 4 seconds, RARE factor = 16, number of averages = 12, field of view (FOV) of 24 mm × 18 mm and 0.9 mm thick slice, and a data matrix of 256 × 192 and 25 interleaved ascending coronal (axial) slices covering the entire brain from the rostral-most extent of the prefrontal cortical region and caudally to the upper brainstem/cerebellum. Saturation bands were applied around the brain to suppress non-brain (muscle) signal.

Functional images were collected using a single-shot spin-echo echo planar imaging (EPI) sequence (26) with the following parameters: TE = 15 ms, TR = 2 seconds, 300 repetitions, FOV = 24 × 18 mm and 0.9 mm thick slice, and a data matrix of 64 × 48 with 25 interleaved ascending coronal slices in the same space as the T2 anatomical. A fat saturation pulse was used to suppress chemical shift artifact and phase correction parameters were acquired during receiver gain adjustment for Nyquist ghost correction. Respiratory rates, isoflurane concentration, body temperature, lighting, and room conditions were kept constant across subjects (18).

### Image pre-processing

All fMRI and anatomical image pairs were reoriented to ensure LPI orientation prior to processing. Binary masks outlining rat brain boundaries were generated in MATLAB using Three-Dimensional Pulsed Coupled Neural Networks (PCNN3D) on the high resolution T2 Turbo-RARE anatomical scans and on fMRI scans (27). N4 bias field correction (28) was used to remove B1 RF field inhomogeneities and reduce FOV intensity variations in T2 anatomical scans prior to alignment to a parcellated rat brain template (29). The binary ‘brain-only’ masks were multiplied by T2 anatomical and fMRI scans to remove voxels outside the brain. The cropped T2 anatomical brain images were aligned to the rat brain T2 template using the FMRIB Software Library (FSL 6.0.3) linear registration tool, FLIRT (30). Linear transform matrices were stored and later applied to functional scans to align these with the same template space.

Resting state processing was carried out using in house UNIX bash scripts that sequentially called tools in Analysis of Functional NeuroImages (AFNI) (31), FSL, and Advanced Normalization Tools (ANTs) (32) to process all images. First, in AFNI we used *3dDespike* to remove time series spikes, *3dTshift* for slice timing correction, and *3dvolreg* for motion and linear drift correction. Cropped fMRI scans were split into 300 individual images and the first in the series was first linearly registered to the corresponding cropped T2 anatomical using FSL FLIRT and then in ANTs we used antsRegistrationSynQuick.sh to warp the fMR image to the same cropped anatomical. Linear and warping matrices were stored and applied to the remaining fMR images in the series. After alignment between the fMRI and corresponding anatomical scans was improved, linear registration matrices transforming fMRI scans to the T2 rat brain template space was applied. Finally, fMRI scans were merged into a single image time series file. In the template space, we regressed white matter and CSF (ventricle) signals, applied bandpass filtering to time series images (0.009 Hz – 0.12 Hz) and spatial smoothing (0.5 mm FWHM), and voxel time series L2 normalization (31).

### Extraction of spontaneously fluctuating fMRI signals for functional connectivity analysis

A total of 144 region of interest (ROI) masks, divided into 72 left and 72 right ROI’s, were included in our analyses. Individual ROI masks were generated using a previously published rat brain parcellation and its accompanying T2 weighted image template (29). To generate the ROI seeds we first produced left and right hemispheric versions of the parcellations, split these into individual left and right image files, extracted center of mass local coordinates for each image, and at these spatial locations we created images with a 0.5-mm radius ROI sphere. We merged 72 left and 72 right hemisphere seeds into a single image file, the coordinates were saved and imported to BrainNet Viewer in MATLAB for 3D visualizations (33) (see below). Seed atlas and coordinate files (and BrainNet viewer files) are all available upon request to the corresponding author. The average principal timeseries vector was extracted from preprocessed fMRI scans with the assistance of ROI mask overlays . This generated 144 individual ROI text files per subject that contained L2-normalized resting state signals as a vector of 300 data points. The timeseries files were used in cross-correlations and in calculations of Pearson r coefficients for every pairwise combinations of ROIs (1dCorrelate in AFNI). The resulting number of pairwise correlations was 10,152 per subject. Correlation coefficients were imported to MATLAB and Fisher’s transform applied to ensure a normal distribution of z values prior to analyses.

### Application of graph theory algorithms to analyze brain wide functional connectivity networks

Functional connectivity graphs were analyzed using Brain Connectivity Toolbox (7). Pairwise correlations were reorganized into symmetrical graphs containing 144 nodes and 10,152 edges, with self-correlations zeroed. Graph edge weights were normalized to a maximum value of 1 (while retaining their normal distribution). To avoid any bias in the calculated network measures, due to inter-subject variations in edge densities, we applied a threshold that maintained an equal edge density across subjects. A percentage of top ranked z values were retained and z values under the set threshold were zeroed. Using this procedure, network measures were calculated for 9 different edge density thresholds ranging from 5-45% (in 5% steps). Following normalization and thresholding, graphs (*G*) were binarized such that *G*_*ij*_ = 1 indicates the existence of a ‘connection’ between any nodes *i* and *j*. For each node, the sum of above-threshold connections in half of the binarized graph provides the *node degree* (total nodal connections). For non-binarized (weighted) graphs, the sum of above-threshold edge weights (z values) provides the *node strength* (strength of total nodal connections). All network calculations were carried out using weighted undirected graphs. Also, for all brain functional network quantifiers we also estimated these for randomly rewired matrices (details below).

### Small world coefficient

To assess network integration and efficiency, we first analyzed the clustering coefficient (*CC*; an index of the average number of connected neighbor pairs *j k* of a node *i* that are connected to each other)(34), characteristic path length (*L*; the lowest average number of edges between any pair of nodes *i* and *j* in graph *G*), and the small world coefficient (σ; which is > 1 for small world networks with *CC* greater than random networks and *L* near random) (6). For *L*, we first substituted matrix weights by their inverse values such that highest z values correspond to shortest distances. Global efficiency was determined as the average inverse of *L*. To calculate σ, we randomly swapped (10 times) all edges in each normalized weighted graph (*G*_*orig*_) to generate a randomized version of the original graph (*G*_*rand*_) that preserved degree and strength distributions (7, 35). *CC* and *L* were then determined for original and randomized graphs and σ calculated as CC_*orig*_/CC_*rand*_ / *L*_*orig*_/*L*_*rand*_ (36).

### Assortativity and rich club index

We next analyzed the tendency of assortative vs dissortative mixing of nodes (37). The assortativity index is a Pearson correlation coefficient comparing node strength values between pairs of edge-connected nodes. Positive r values indicate connectivity between pairs of nodes with similar strengths(e.g., high strength nodes pairs with high and low with low), while negative r values indicate cross-pairings between low and high strength nodes. We also generated and analyzed weighted rich club coefficient curves (*Φ*) at an edge density of 10% (38). Nodes belonging to a rich club subnetwork have an above-chance tendency to tightly connect with each other and share the bulk of connections within the network (38, 39). The approach creates subgraphs containing nodes with strength values at a predetermined degree value, *k*. For each k-level subgraph, *Φ*_*w*_ is calculated as the ratio of the sum of edge weights that connect nodes of degree k or higher (*W*_*>k*_) to the total possible number of summed edge weights(39):

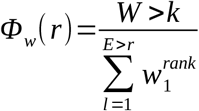

### Modularity and Node Cartography

Functional connectivity has been previously reported to follow a modular organization in which subgroups of nodes show a greater degree of intra-group functional connectivity than inter-group connectivity (40). To assess the effect of context exposure and social interaction on this segregated pattern of activity we calculated a probabilistic version of the modularity index (41). The agglomerative procedure starts with a random grouping of nodes and iteratively moves nodes into groups which maximize a modularity index, *Q*. The final module assignments for each node (e.g., a community affiliation vector) was taken as the median of 1000 iterations of the modularity maximization procedure. The role each node plays within their assigned module can vary according to their within-module degree (relative to other nodes in the same module) and their level of participation in connections with nodes in other modules. To investigate the role of intra-module nodes, we analyzed the within-module degree z-score (*z*) and the participation coefficient (*P*) (42). A node *i* in module *m*_*i*_ may play a central role if it has a within-module node degree (*d*_*i*_) greater than the average node degree (*δ*) for that module,

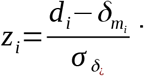

The difference between individual intra-module node degree and average intra-module node degree is variance-normalized to produce a z-score which allows the assignment of connector hubs at *z*_*i*_>2.5 (*z*_*i*_>1 in functional connectivity networks) (42, 43). The participation coefficient is calculated as:

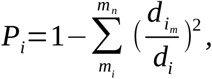

where *d*_*im*_ is the within-module node degree and *d*_*i*_ is the overall network degree for a node *I (42)*. Within-module nodes with different levels of participation in connections with nodes in other modules were classified as follows: *P*<0.05 are ultra-peripheral, 0.05<*P*<0.62 are peripheral nodes, 0.62<*P*<0.80 are non-hub connectors, and *P*>0.80 are non-hub kinless (42).

### Centrality and network robustness

Functional connectivity networks are thought to contain high degree ‘hub’ nodes that either share connections with a large proportion of nodes, connect with other high degree nodes, and/or have high transit of connections between pairs of nodes in a network (44). To measure the effect of context exposure and social interaction on putative hub node configurations we calculated betweenness and eigenvector centrality. A node *i* has high betweenness centrality if it lies within many shortest paths between any nodes *j k* (relative to all shortest paths between *j k*) (44). Eigenvector centrality considers not only the number of connections node *i* possesses (quantity) but places greater weight on how many of these are with high strength nodes (quality) (40).

In addition, we tested network robustness at an edge density of 10% using a random and preferential node removal strategy, as previously proposed (5, 45). For random node removal (‘random failure’), we used a random permutation approach that iteratively removed individual nodes, and at each removal step calculated the size of the largest component. The size of the largest connected component (*S*) was averaged over 1000 iterations and proportionally scaled between 0 and 1. For preferential node removal (‘targeted attack’), nodes are first ranked from highest to lowest values for node degree, strength, betweenness centrality, or eigenvector centrality, and then removed in a step wise manner while calculating *S* at each step. We compared the results of random and targeted attack in rat fMRI networks with synthetically generated random and ring lattice networks with the same number of nodes and edge densities.

### Functional connectivity network visualizations

Functional connectivity networks were visualized in BrainNet viewer (33). Center coordinates for each ROI were generated in BrainNet viewer based on parcellations on a previously published rat brain template (29). A 3D whole brain surface mesh file of the rat brain template (in *.byu format) was generated using an image binarization command in FSL (*fslmaths*) (30) and mesh construction in ITKSNAP (46). Several 3D brain maps were generated. A first series of connectivity strength maps were produced with the size of nodes (spheres) weighted by node strength and the edges (lines connecting node pairs) weighted by the Pearson coefficient between pairs of nodes (lower bounds threshold set at z=0.15). Additional maps were generated with node color and edges representing the module assignment (e.g., community affiliation vector) and node size weighted by either the modularity index, local efficiency, betweenness, or eigenvector centrality scores.

### Statistical analyses

Statistical analyses of network measures and data plotting were carried out in MATLAB. Unless otherwise indicated in the results section and figure captions, analysis of variance (ANOVA) was used with pre-scan manipulations as independent variables and Tukey-Kramer post hoc test for multiple comparisons (p<0.05).

### Functional MRI reanalysis: comparing results with an alternative pre-scan intervention

We reanalyzed a subset of control scans that were previously collected as part of a different set of studies. This additional analysis was carried out to determine the degree of specificity of the present results by assessing whether a different intervention prior to imaging exerted a similar or different effect on resting state functional connectivity and connectome quantifers. We determined the effect of a single saline injection administered either 1 hour (n=6) or 24 hours (n=10) prior to fMRI scanning (23) and compared these two groups to a naive group of rats scanned with no pre-scan intervention (n=8) (47). Saline injections were applied over a lower abdominal quadrant using sterile packaged tuberculine (1cc) syringes and 26 gauge needles, at a 45 degree angle into the intraperitoneal layer. Following injections, rats were returned to their home cages and scanned according to previously reported methods (23, 47). All image processing, network calculations and statistical analyses and plotting were carried out as described in the previous sections.

## RESULTS

**Figure 2.**
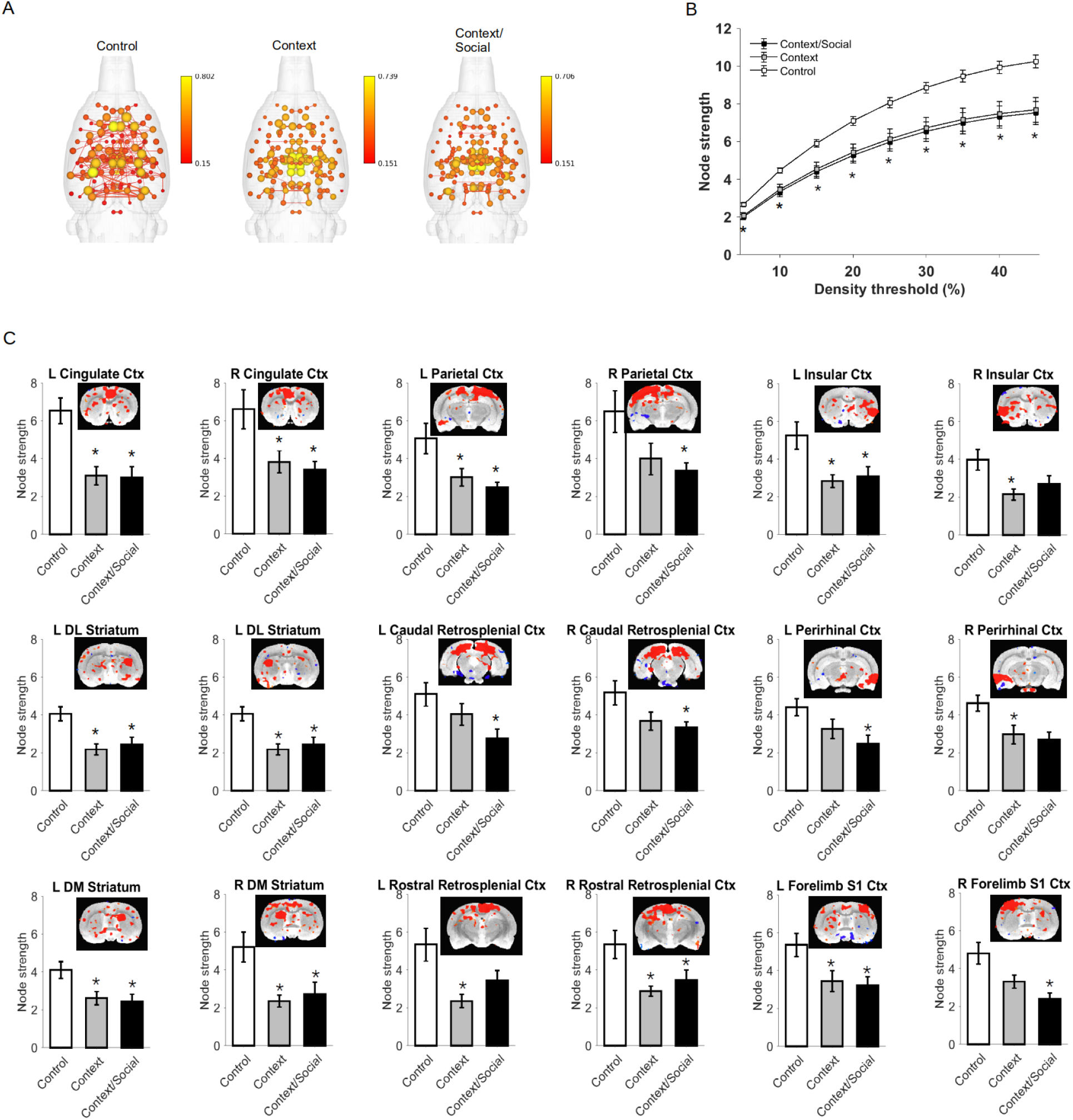
Presentation of a familiar context prior to imaging resulted in significantly reduced node strength in cortical regions involved in memory. A) Connectome maps with anatomically-placed nodes scaled by strength (edges are normalized Pearson’s r thresholded at 0.15).Scale bar represents edge weight values. B) Global node strength at different graph edge densities. C) Local node strength was reduced bilaterally in cortical regions and striatum. Functional connectivity maps at level seed location and overlaid onto T2 template are shown above bar graphs. Data are mean ± standard error. *Significantly different from control (ANOVA with Tukey-Kramer post hoc test). Abbreviations: DM, dorsomedial; DL, dorsolateral; L, left; R, right; Ctx, cortex; S1, primary somatosensory.

### Context exposure significantly reduced network strength

We observed a significant reduction in node strength at all graph densities following context exposure (5%-45%: F_2,27_>7.6, p<0.005) but no distinct effect of the social interaction within the same context (Figure 2B). Post hoc tests indicated significant reductions in context and context/social groups relative to controls (p<0.05). No differences between context and context/social groups were observed. Differences in node strength were also observed at the nodal level in left and right cingulate (left: F_2,27_= 11.7, p = 0.0002; right: F_2,27_= 5.3, p = 0.01), parietal (left: F_2,27_= 5.4, p = 0.01; right: F_2,27_= 3.6, p = 0.04), insular (left: F_2,27_= 5.7, p = 0.008; right: F_2,27_= 4.6, p = 0.02), caudal (left: F_2,27_= 4.2, p = 0.03; right: F_2,27_= 3.8, p = 0.04) and rostral retrosplenial (left: F_2,27_= 5.7, p = 0.008; right: F_2,27_= 5.3, p = 0.01), and perirhinal cortices (left: F_2,27_= 4.2, p = 0.02; right: F_2,27_= 5.7, p = 0.008), forelimb (left: F_2,27_= 4.6, p = 0.02; right: F_2,27_= 7.4, p = 0.003) and jaw somatosensory regions (left: F_2,27_= 10.8, p = 0.0004; right: F_2,27_= 3.9, p = 0.03), secondary visual cortex (left: F_2,27_= 3.8, p = 0.04; right: F_2,27_= 3.8, p = 0.03), and dorsomedial (left: F_2,27_= 5.2, p = 0.01; right: F_2,27_= 6.5, p = 0.005) and dorsolateral striatum (left: F_2,27_= 8.6, p = 0.001; right: F_2,27_= 6.2, p = 0.006). Post hoc results summarized in Figure 2C indicated that either context, context/social or both reduced node strength bilaterally in these nodes (p<0.05). In addition, we observed reductions in node strength in other unilateral regions. These included in the right hemisphere of central amygdala (F_2,27_=5.2, p=0.01), primary visual (F_2,27_=6.9, p=0.004) and auditory cortex (F_2,27_=3.6, p=0.04) secondary somatosensory cortex (F_2,27_=5.8, p=0.008), dorsal hippocampal CA3 (F_2,27_=3.6, p=0.04), periaqueductal grey (F_2,27_=3.7, p=0.04), anterior thalamic nucleus (F_2,27_=8.1, p=0.002). In the left hemisphere we observed reduced node strength in ventral orbital (F_2,27_=4.8, p=0.02), primary (F_2,27_=7.0, p=0.004) and secondary motor cortices (F_2,27_=3.4, p=0.047) and barrel field (F_2,27_=5.1, p =0.01), hindlimb (F_2,27_=6.4, p=0.005), shoulder (F_2,27_=11.4, p=0.0002) and trunk (F_2,27_=6.5, p=0.005) somatosensory regions, lateral septal nucleus (F_2,27_=4.5, p=0.02), ventrolateral (F_2,27_=5.6, p=0.009) and ventromedial striatum (F_2,27_=9.1, p=0.0009), and lateral geniculate (F_2,27_=3.9, p=0.03).

**Figure 3.**
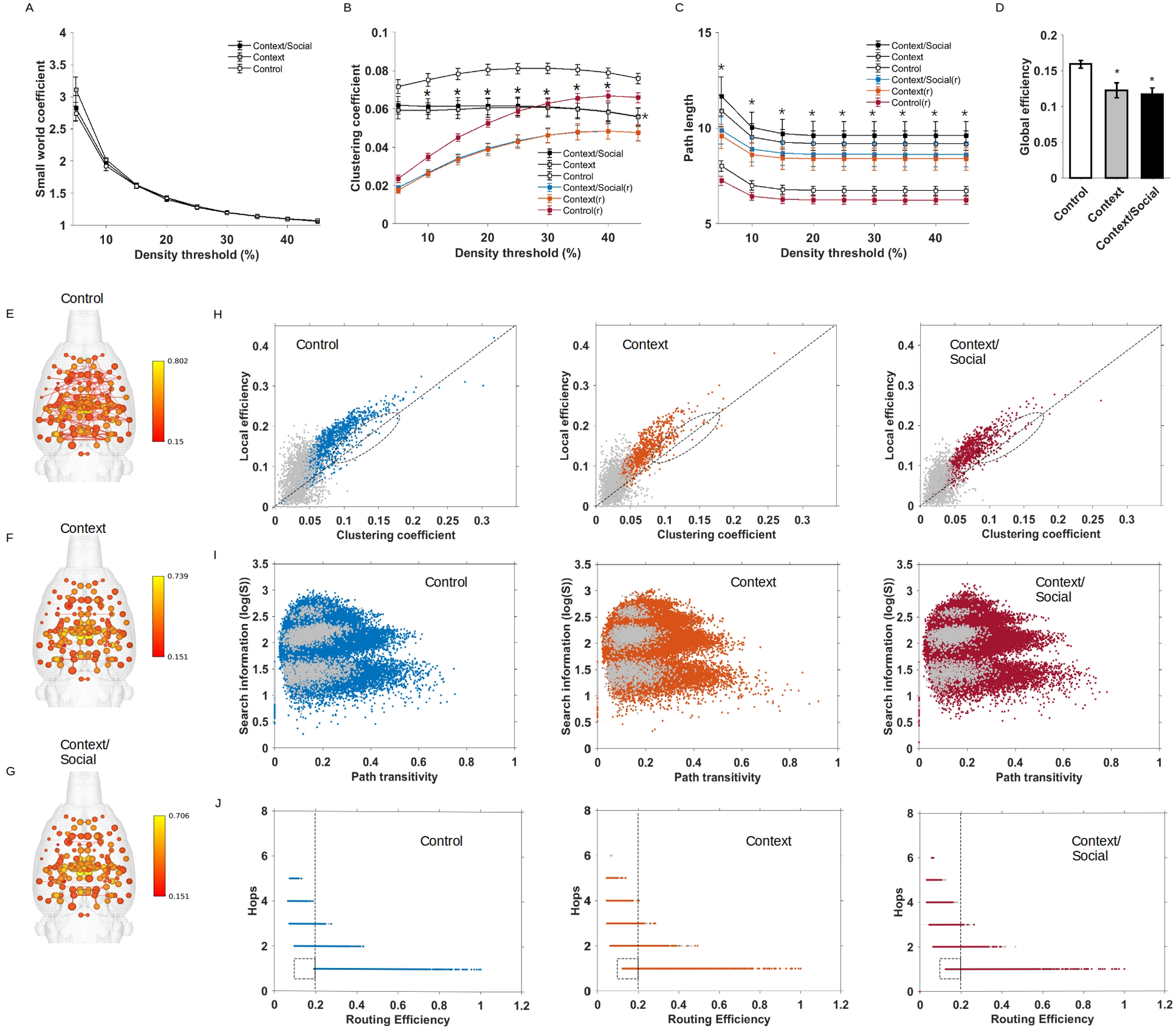
Global efficiency is reduced by context exposure. A) Small world index at various densities. B) Clustering coefficient at various densities. C) Characteristic path length at various densities. D) Global efficiency. E-G) 3D connectome maps with nodes scaled by local efficiency in control (E), context exposed (F), and context/social exposed (G). Sale bar represent edge weights values (threshold r=0.15). H) Distribution of local efficiency vs. clustering coefficient values for all nodes and all subjects in control, context and context/social exposed rats. I) Distribution of search information vs path transitivity values for all edges and all subjects in control, context and context/social exposed rats. J) Distribution of number of steps or hops between node pairs vs routing efficiency values for all edges and all subjects in control, context and context/social exposed rats. In B-C, blue, orange, red line plots are for randomized (r) versions of functional connectivity matrices. In H-J, gray color dots represent data points for nodes or edges of randomized graphs. Data are mean ± standard error. *Significantly different from control (ANOVA with Tukey-Kramer post hoc test).

### Context exposure significantly reduced global network efficiency and clustering

No differences in the small world index was observed between the groups. However, we observed significant differences in the global weighted values for the clustering coefficient (10%-45%: F2,27>5.4, p<0.01) and average characteristic path length (5%-45%: F2,27>7.1, p<0.005). Post hoc results reported in Figure 3B-C indicate that context and context/social groups had lower clustering and higher path length values relative to controls (p<0.05). As expected from the low global clustering and long average path lengths, global efficiency values were also significantly lower in context and context/social groups relative to controls (F2,27=7.6, p=0.002) (Figure 3D). Compared to global efficiency, and based on the size of nodes in 3D connectome maps shown in Figure 3E-G, average values for local efficiency were similarly yet less robustly reduced at the individual node level (Figure E-G). Analysis of local efficiency values across all nodes in all rats (as a function of local clustering coefficient values) suggested relatively minor reductions local efficiency in context and context/social groups compared to controls (Figure 3H). This suggested that while context exposure produced robust reductions in global efficiency associated with reduced node strength, clustering, and increased path lengths, this globally reduced efficiency was not widespread in all nodes. Based on this possibility, we hypothesized that the particular arrangement of edges connecting nodes may be important in preserving local efficiency. If so, there would be relative few differences in edge quantifiers that support reduced local efficiency. To investigate this, we calculated search information, which estimates the amount of information (bits) needed to randomly trace shortest paths between node pairs (48). We plotted search information as a function of path transitivity, which estimates the density of detours lying between node pairs (48). This analysis indicated no differences in terms of accessibility or degree of difficultly in randomly finding shortest paths between node pairs in control, context, and context/social exposure rats (Figure 3I). We next plotted the number of edges (termed ‘hops’) lying along shortest paths between a pair of nodes as a function of routing efficiency (the average of inverse shortest path length)(49–51). As anticipated, differences in efficiency were subtle and limited to single ‘hops’ which had slightly lower routing efficiency in context and context/social groups relative to controls (Figure 3J).

Finally, we note that in randomized functional connectivity matrices the estimated values for clustering coefficient and characteristic path lengths were lower than in original matrices, although these preserved node strength differences between the groups (Figure 3). Also, while global efficiency values were greater in randomized versus original matrices (data not shown), local efficiency and path transitivity values were much lower in rewired networks (Figure 3H-I).

**Figure 4.**
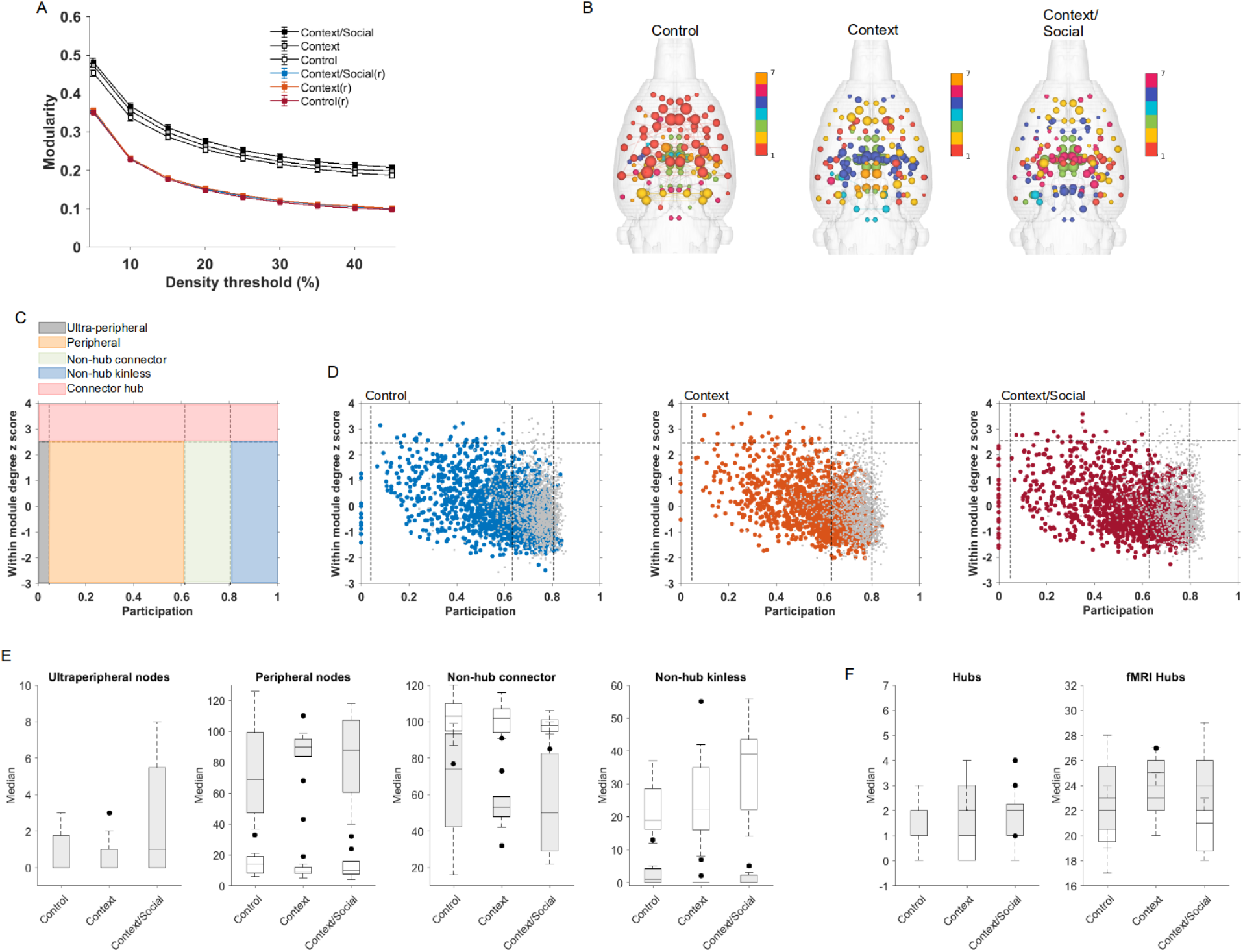
Context exposure did not alter modular organization of nodes. A) Modularity index at various densities. B) 3D connectome maps with nodes scaled by modularity index Q in control, context exposed, and context/social exposed. Scale bar color represent community affiliation of nodes. C) Nodal assignment chart according to (42). D) Scatter plots of within module degree z score versus participation coefficients per node for all subjects in control (blue), context (orange) and context/social (red) exposed rats. Hashed lines represent limits for specific ranges for cartographic assignments per node according to (44). E) Nodal role assignments according to participation coefficient values. F) Nodal assignments according to within module degree z scores (hubs > 2.5; fMRI hubs > 1 (43)). In A, blue, orange, red line plots are for randomized (r) versions of functional connectivity matrices. In D, gray color dots represent data points for nodes or edges of randomized graphs. In E-F, data are median ± 95% confidence intervals (box-whisker plots, with black dots representing outliers). Gray boxes are for real fMRI networks and empty are for randomized versions of the same networks. Data are mean ± standard error. *Significantly different from control (ANOVA with Tukey-Kramer post hoc test).

### No significant effects of context or social exposure on modularity or node assignments

No effect of stimuli on global modularity was observed (Figure 4A). The number of total affiliations or groups did not vary between the 3 groups, although on average an apparent difference in the assignment of nodes to modules was observed in group-averaged 3D connectome maps (Figure 4B). To further investigate the role of nodes within the identified modules, we used an approach published by (42) and updated for functional MRI data sets by Meunier *et al* (43)(Figure 4C-F). Based on the within-module degree z-score (at a threshold of 10%) and using a criterion score of z >2.5 proposed by Guimares and Amaral (42), we observed very few connector hubs across all subjects in the 3 groups. Using a criterion score of z >1 for functional connectivity networks of proposed by Meunier *et al* (43), we observed a median value of 20-25 connector hubs across rat functional networks (Figure 4F). Guided by the participation coefficient parcellation, we observed that a large proportion of nodes across all subjects and groups were peripheral nodes followed in quantity by non-hub connectors (Figure 4D-E). To a lesser degree, there were ultra-peripheral nodes and virtually no non-hub kinless. There was no effect of cage or social exposures on any of these nodal role assignments.

Finally, modularity index values were significantly lower in randomly rewired functional networks compared to original versions. Also, there were lower fMRI hubs in randomly rewired versus original networks and the distribution of participation coefficient values indicated that rewired networks only had nodes with roles as non-hub kinless and non-hub connectors.

**Figure 5.**
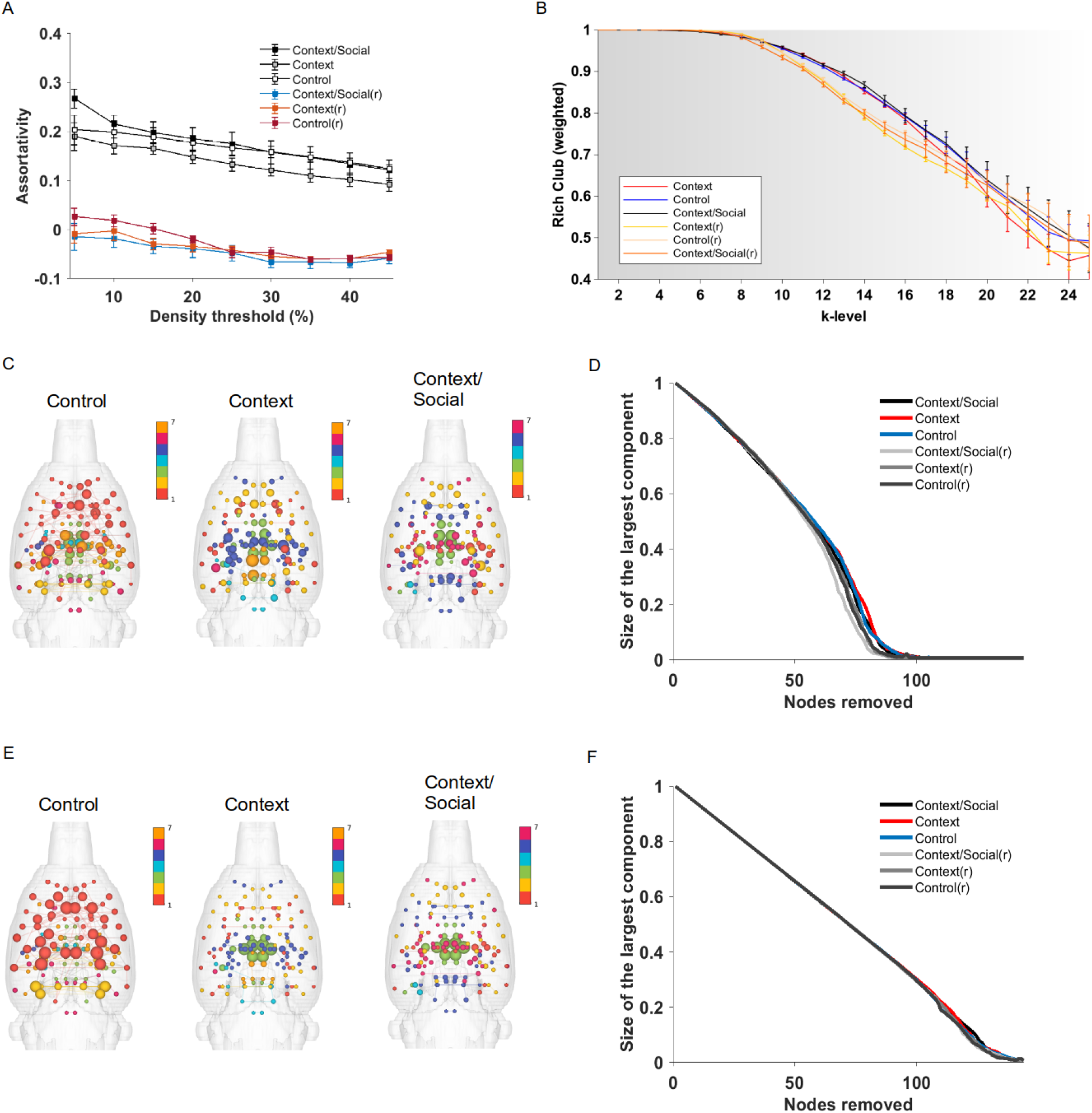
Context exposure did not alter functional connectivity between strongly weighted nodes and did not alter network robustness. A) Assortative mixing at various graph densities. B) Rich club curve for a 10% graph density. C) 3D connectome maps with nodes scaled by betweenness centrality ranking in control, context exposed, and context/social exposed. Scale bar color represent community affiliation of nodes. D) Size of the largest component as a function of node removal according to descending betweenness centrality ranking. E) 3D connectome maps with nodes scaled by eigenvector centrality ranking in control, context exposed, and context/social exposed. Scale bar color represent community affiliation of nodes. F) Size of the largest component as a function of node removal according to descending eigenvector centrality ranking. Data in A-B are mean ± standard error. In A, blue, orange, red line plots are for randomized (r) versions of functional connectivity matrices. In B, yellow-orange line plots are for randomized (r) data sets. In D and F, gray scale colored curves are for randomized (r) data sets.

### No significant effects of context or social exposure on rich club organization or assortative mixing

No effects of the stimuli was observed on assortative mixing or the rich club index (Figure 5A-B). Assortativity index values were lower in randomized networks versus original versions. These were around zero, indicating randomized patterns of connectivity relative to the original versions containing greater number of connections between high strength nodes. Similarly, rich club index values at a specific range of k-level values between 10-16 were lower in the randomized versus original functional networks.

### No significant effects of context or context/social stimulus exposure on functional network robustness

Preferential or targeted removal of highly connected hub nodes can cause a sharp reduction in the size of the largest component, *S,* and reveal the critical point at which the network breaks into much smaller unconnected components (40, 52–54). Sharper or faster reductions in *S* as a function of the fraction of nodes present is indicative of a high reliance on hub nodes, which are central to overall network robustness (45). We carried out the targeted node removal process based on descending rank of either node strength, node degree, betweenness centrality, or eigenvector centrality scores (Figure 5C-F). No effect of context or context/social stimulus exposure was observed on network robustness. Results for betweenness centrality and eigenvector centrality are shown in Figure 5C-F.

**Figure 6.**
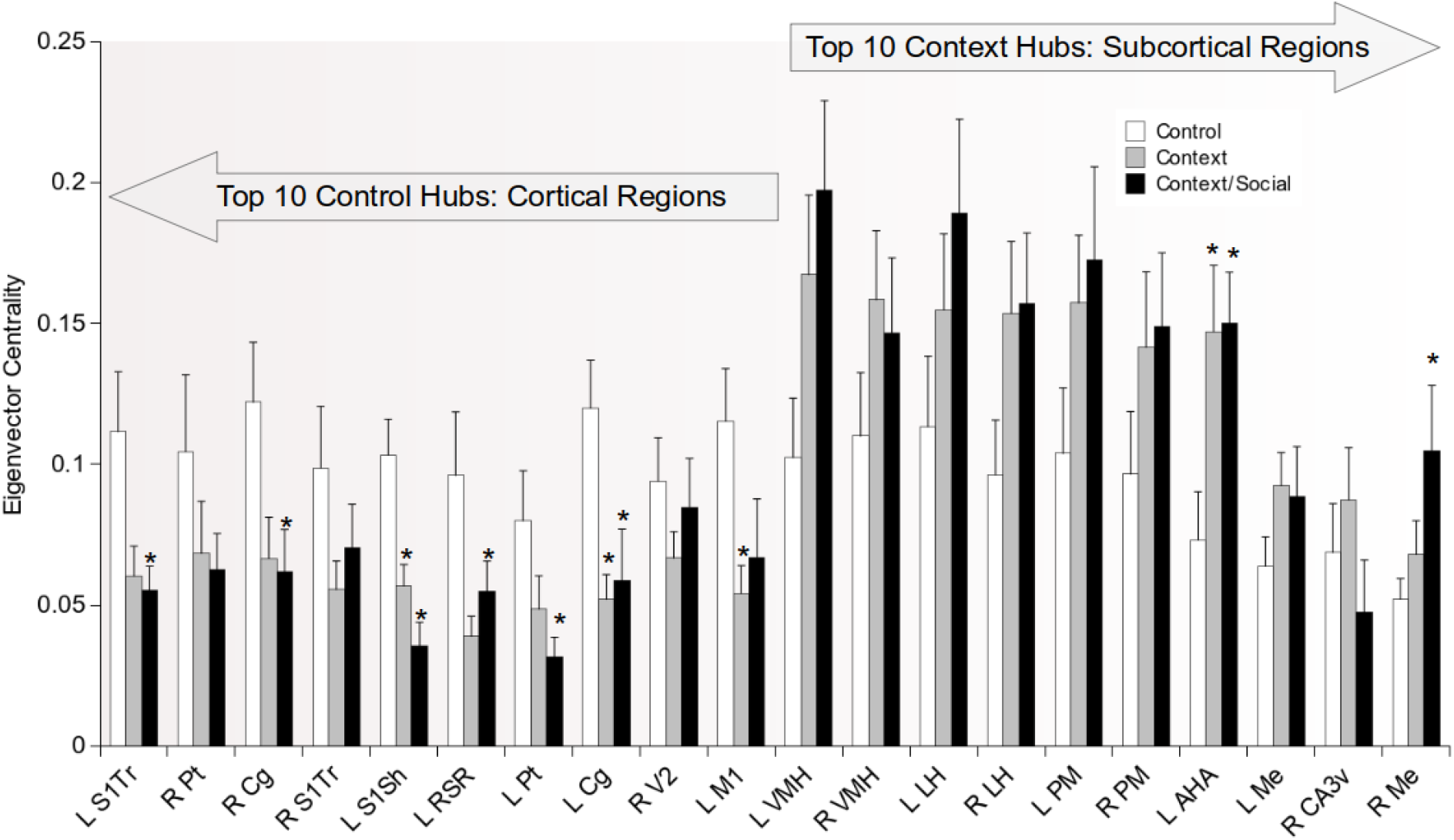
Context and context/social exposure resulted in a switch in high eigenvector centrality node locations from cortex to subcortical regions. Eigenvector centrality rankings of top 10 nodes in control rats and in context exposed rats (arrows above bar graphs indicate top 10 nodes in control vs context exposed rats). Data are mean ± standard error. *Significantly different from control (ANOVA with Tukey-Kramer post hoc test). Abbreviations: L, left; R, right; S1, primary somatosensory; Tr, trunk; Sh, shoulder; Cg, cingulate, Pt, parietal; RSR, rostral retroplenium; V2, secondary visual cortex; M1, primary motor cortex; VMH, ventromedial hypothalamus; LH, lateral hypothalamus; PM, posteromedial thalamus; AHA, anterior hypothalamuc area; Me, medial amygdala; CA3v, ventral CA3 of hippocampus.

### Context and context/social exposure resulted in a switch in high eigenvector centrality node locations from cortex to subcortical regions

The group-averaged 3D connectome maps in Figure 5E suggested that exposure to a familiar context resulted in a significant reduction in node eigenvector centrality values. We sorted average node eigenvector centrality values from highest to lowest and selected the top 10 in each group. Results are summarized in Figure 6. The top 10 nodes with highest eigenvector centrality values (which we refer to here as hubs) in control rats were cortical. These included cortical areas observed to have reduced node strength in response to familiar context exposure. Many of these same regions showed a significant reduction in eigenvector centrality and this effect was observed in the trunk (F_2,27_=4.2, p=0.03) and shoulder regions of primary somatosensory cortex (left: F_2,27_=11.9, p=0.0009), and parietal (left: F_2,27_=3.2, p=0.05), cingulate (left: F_2,27_=6.3, p=0.006; right: F_2,27_=3.7, p=0.04), retrosplenial (left: F_2,27_=3.6, p=0.04) and primary motor cortical regions (left: F_2,27_=3.8, p=0.04). The group-averaged 3D connectome maps for the familiar context exposure groups in Figure 5E also suggest that cortical hubs are significantly suppressed. Indeed, these maps suggest that nodes in deep subcortical areas have the highest eigenvector values. Upon sorting nodes from highest to lowest eigenvector values in these groups, we observed that the top 10 nodes were subcortical - largely hypothalamic. Among these regions, the anterior hypothalamic area had eigenvector centrality values that were greater in context exposed compared to control animals (left: F_2,27_=5.1, p=0.01).

**Figure 7.**
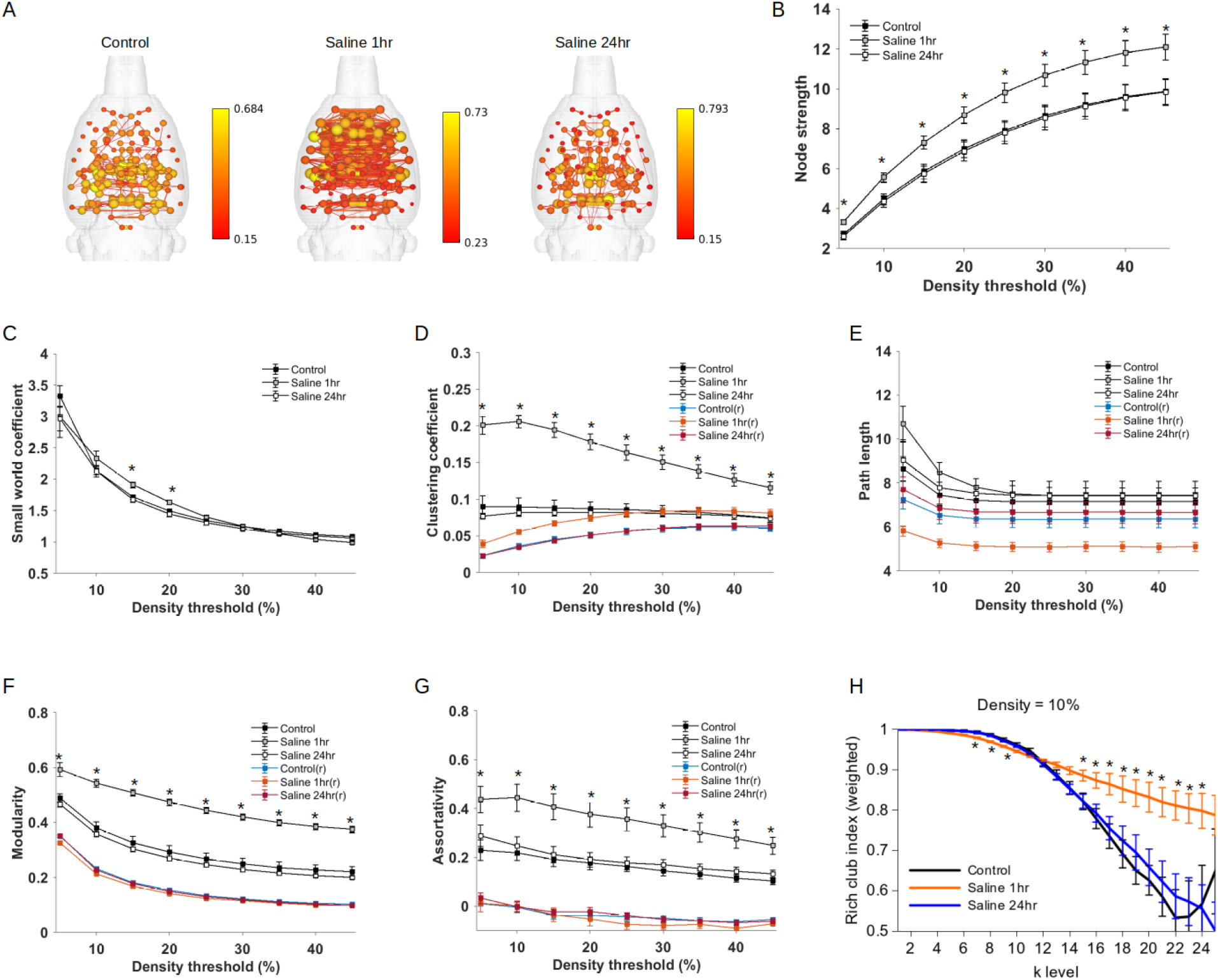
Functional connectome changes in response to an arousing stimulus administered to rats at the same temporal delay prior to imaging as the context and social/context exposures. The arousing stimulus was a manual handling procedure during administration of an intraperitoneal injection of sterile physiological saline solution. A) Connectome maps with anatomically-placed nodes scaled by strength (edges are normalized Pearson’s r thresholded at 0.15).Scale bar represents edge weight values. B) Global node strength at various graph densities. C) Small world index at various graph densities. D) Clustering coefficient at various graph densities. E) Characteristic path length at various graph densities. F) Modularity at various graph densities. G) Assortativity index at various graph densities. H) Rich club curve at a 10% graph edge density. In B-G, blue, orange, red line plots are for randomized (r) versions of functional connectivity matrices. Data are mean ± standard error. *Significantly different from control (ANOVA with Tukey-Kramer post hoc test).

### An arousing saline injection stimulus prior to imaging produced network topological changes that are diametrically distinct from familiar context exposure

We reanalyzed an MRI data set in which rats were scanned either 1 or 24 hours after an unexpected and unfamiliar arousing stimulus, an intraperitoneal injection of sterile saline solution. These groups were compared to a naive control group of age, sex, weight, vendor and imaging protocol matched rats that were scanned directly upon arrival at the imaging facility and did not receive any pre-scan intervention. This ancillary study was conducted to determine if a general arousing stimulus could produce similar topological changes in functional connectivity networks in rats. First, values for node strength, small world index, clustering coefficient, characteristic path length, modularity, assortativity and rich club index for control rats imaged at 4.7 Tesla were almost identical to those calculated for functional connectivity data of rats imaged at 11.1 Tesla (control rat data in Figure 7 versus Figures 2–5). 3D connectome maps in Figure 7A show stronger overall connectivity and average node strength in rats injected 1 hour prior to image compared to those injected 24 hours earlier, or the naive control group. Node strength was much greater in 1 hour vs 24 hour and control rats across several network density thresholds (5%-30%: F_2,21_>3.6, p<0.05)(Figure 7B). The small world index was slightly greater in 1 hour injected rats versus 24 hour and control animals, although this was observed only at 15-20% density thresholds (15%: F_2,21_=8.4, p=0.002; 20%: F_2,21_=8.0, p=0.003) (Figure 7C). In addition, the injection 1 hour pre-scan resulted in a significantly greater clustering coeffiicient (5%-45%: F_2,21_>14, p<0.0001), greater modularity (5%-45%: F_2,21_>16, p<0.0001), assortativity (5%-45%: F_2,21_>4.5, p<0.05) and rich club organization (k-level 6-8 F_2,21_=6.9, p=0.005 and 14-24 F_2,21_=6.7, p=0.006) than rats injected 24 hours earlier or naive control animals (Figure 7D-H). Interestingly, the injection 1 hour prior to scanning did not alter characteristic path length. In sum, the response to a general arousing stimulus such as an intraperitoneal injection administered 10 minutes prior to sedation and 50-60 minutes prior to fMRI scanning caused an almost lockstep increase in functional connectivity network parameters. This is in stark contrast with the network changes caused by exposure to a familiar contextual stimulus.

**Figure 8.**
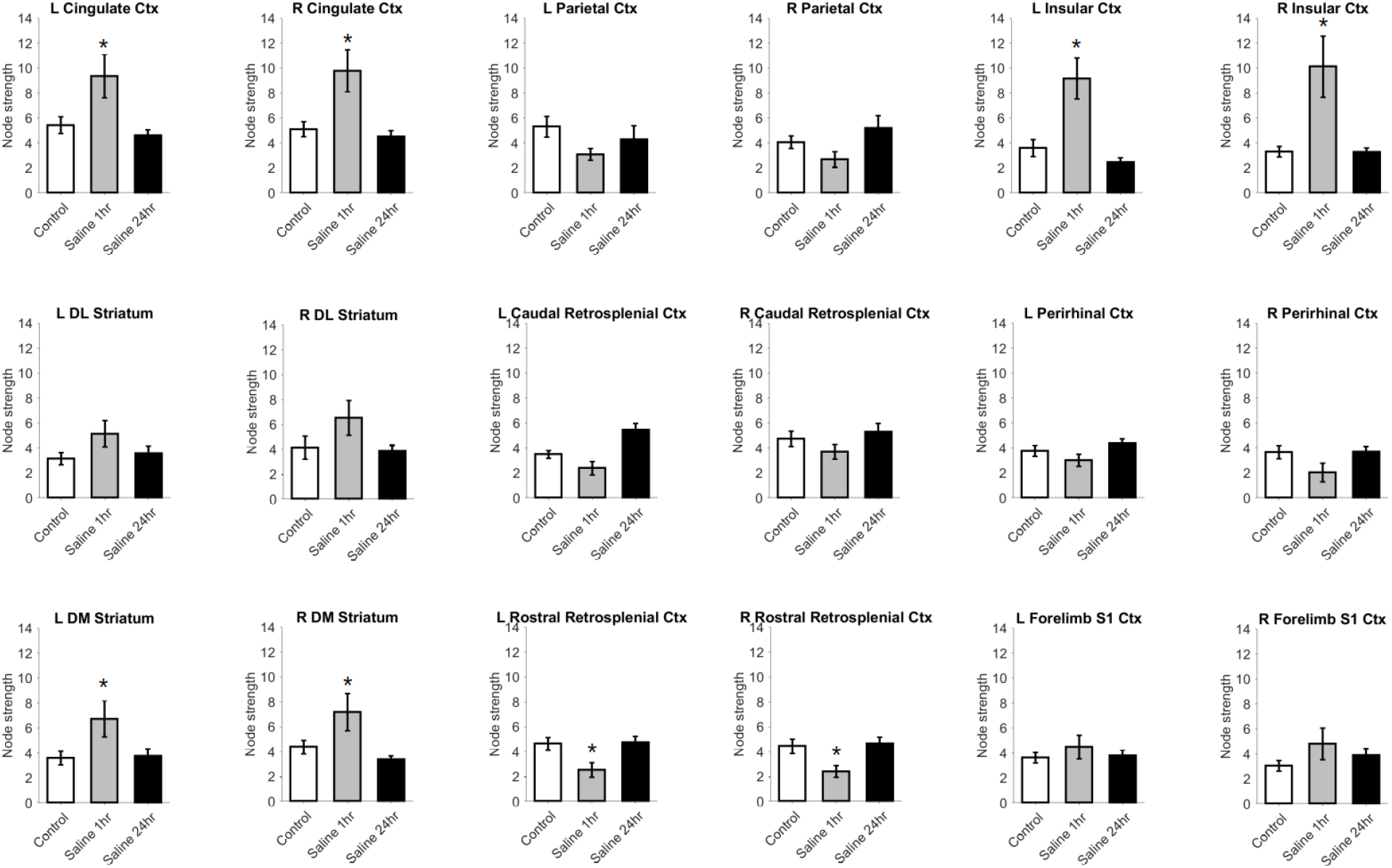
Differential effects of the arousing stimulus on local node strength in regions also assessed in response to context and social/context exposures (Figure 2). Data are mean ± standard error. *Significantly different from control (ANOVA with Tukey-Kramer post hoc test). Abbreviations: DM, dorsomedial; DL, dorsolateral; L, left; R, right; Ctx, cortex; S1, primary somatosensory.

To further determine if the general arousing injection procedure produced network changes that are distinct from context exposure, we analyzed node strength in regions that had reduced node strength with familiar context exposure (Figure 8). This additional analysis revealed differences in local connectivity strength between the two studies. The 1 hour injection increased node strength in cingulate (left: F_2,21_=6.4, p=0.007; right: F_2,21_=9.1, p=0.001) and insular cortices (left: F_2,21_=14.1, p=0.0001; right: F_2,21_=10.9, p=0.0005) and dorsomedial striatum (left: F_2,21_=4.4, p=0.03; right: F_2,21_=5.5, p=0.01), and reduced node strength in the rostral retrosplenial cortex (left: F_2,21_=4.6, p=0.02; right: F_2,21_=4.3, p=0.03), but had no effect on any other region that were responsive to presentation of a familiar context (Figure 8). In addition, the injection procedure significantly increased node strength in a different set of structures involved in regulation or modulation of affective responses. These included the anterior (left: F_2,21_=6.5, p=0.006; right: F_2,21_=5.1, p=0.02), basal (left: F_2,21_=15.7, p<0.0001; right: F_2,21_=13.3, p=0.0002), central (left: F_2,21_=4.8, p=0.02; right: F_2,21_=14.3, p=0.0001), lateral (left: F_2,21_=11.2, p=0.0005; right: F_2,21_=14.0, p=0.0001), medial (left: F_2,21_=9.0, p=0.002; right: F_2,21_=16.8, p<0.0001) amygdala nodes and bed nucleus of stria terminalis (left: F_2,21_=5.6, p=0.01; right: F_2,21_=4.7, p=0.02), prelimbic cortex (left: F_2,21_=3.7, p=0.04; right: F_2,21_=10.5, p=0.0006), primary (left: F_2,21_=8.4, p=0.002; right: F_2,21_=11.8, p=0.0003) and secondary motor cortices (left: F_2,21_=4.0, p=0.03; right: F_2,21_=3.6, p=0.045), caudal piriform cortex (left: F_2,21_=5.1, p=0.02; right: F_2,21_=8.9, p=0.002), lateral septal nucleus (left: F_2,21_=10.6, p=0.0006; right: F_2,21_=12.5, p=0.0002), medial preoptic area (left: F_2,21_=7.8, p=0.002; right: F_2,21_=6.7, p=0.005), central grey (left: F_2,21_=8.5, p=0.002; right: F_2,21_=7.3, p=0.003), nucleus accumbens (left: F_2,21_=6.5, p=0.006; right: F_2,21_=7.2, p=0.004), ventromedial striatum (left: F_2,21_=15.1, p<0.0001; right: F_2,21_=29.6, p<0.00001), ventral pallidum (left: F_2,21_=13.1, p=0.0002; right: F_2,21_=7.4, p=0.004).

## DISCUSSION

The data indicate that exposure to a familiar context resulted in a significant reduction in connectivity strength in rat brain. Analyses of 72 bilateral brain regions further indicated that node strength values were reduced in areas of the cortex known for their roles in long-term memory. The regions included left and right cingulate, parietal, insular, perirhinal, retrosplenial, and primary somatosensory cortices, and dorsal striatum. In addition to changes in node strength, we observed a significant reduction in weighted measures of clustering and efficiency. This suggests that because of the overall reduction in node strength there is a significant loss of strongly weighted triad structures (weighted clusters) in rats exposed to a familiar context, and this in turn is associated with a reduced potential for information transfer. Finally, while in control rats the top 10 nodes with highest eigenvector centrality values were cortical, in rats exposed to a familiar context the top 10 hub nodes were subcortical. The top 10 hub regions in controls were the same structures that showed reduced connectivity strength in context-exposed rats. Thus, exposure to a familiar context modified neural interactions in such a way that suppressed synchronous cortical hubs, which resurfaced in hypothalamic regions. Similar or distinct changes in network quantifiers in response to the unfamiliar social stimulus were not observed. In general, we conclude that the neural processing of a recognized or familiar stimulus exerts a greater influence in reorganizing network topology than a novel (social) stimulus. We propose that this could be due to the activation of reconsolidation mechanisms in response to the familiar stimulus. Reconsolidation may involve changes in the strengthening of synaptic activity to maintain a degree of fidelity of memory (24, 55, 56). Neural activity in response to unfamiliar stimuli could potentially interfere with reconsolidation mechanisms by adding noise or unrelated neuronal activity to pre-established functional network interactions.

Among unanswered questions in this study, two stand out as critical to address in future studies. The *first* relates to neuronal activity mechanisms underlying the observed reductions in cortical node strength, efficiency, clustering, and cortical hubness. A *second* question relates to the lack of effect of the unfamiliar social stimulus relative to familiar context exposure (57). The first question could be addressed in future studies using local field potential recordings in multiple cortical regions of rats tested under similar experimental conditions as our present neuroimaging study. It is well established that the frequency bandwidth of spontaneous BOLD signal oscillations falls within previously characterized infraslow frequencies (<0.5) that cover a large expanse of rat cortex (58, 59). Studies in primate visual cortex have also shown that normalized spectral frequencies in the band limited power for gamma (40-100Hz) share a high degree of statistical dependency (mutual information) with the fMRI BOLD signal while alpha (8-12 Hz), beta (18-30 Hz), and higher frequency (0.9-3kHz) multi-unit activity much less so (60). Changes in BOLD signal amplitudes scale in a step wise manner with changes in gamma power (60). Thus, somatodendritic processing at band limited power in gamma and infraslow ranges might correspond with synchronous BOLD oscillations underlying functional connectivity. Alterations in gamma band power for a particular frequency might indicate large scale changes in neural processing, which might produce asynchronous cortical BOLD activity. In other words, a switch from gamma to another band limited power spectral frequency, i.e., theta, involved in the processing of information related to a familiar context might underlie reduced connectivity strength in cortical regions. However, it is important to note that this viewpoint is incomplete and research on the large synaptic activity underlying BOLD signal fluctuations is ongoing (61). Although band limited power for gamma shows strong statistical dependency with BOLD, using a sliding window approach others have shown that the relationship between spontaneous fluctuating BOLD signals and underlying neural signals is more complex (62). Using this approach, strong correlations with spontaneous BOLD activity are see in gamma, theta and beta bands (62). Work in human subjects have also reported a closer relationship between alpha and regions of the salience and dorsal attention networks, and theta and frontal parietal and default mode networks (63). Thus, the relationship between BOLD and underlying large scale synaptic activity may be behavior, cognition, affective state-specific.

Regarding the *second* open question stated above, there have been prior reports suggesting a strong impact of familiarity to an environmental context in habituation (learning) processes involved in social interaction and recognition in rats (57). Behavioral changes observed in response to novel social interactions and recognition may depend on familiarity with the associated context (59). The observed reductions in strength, clustering, efficiency and cortical hubs following familiar context exposure might correspond to reconsolidation processes such as the modifications of synaptic strength in relation to inter-regional communication between distributed cortical neuronal populations (63). To maintain the fidelity of synaptic communication related to an already experienced contextual environment, network topology transiently moves to a less efficiency state as a way of minimizing novel-stimulus interference. Reductions in network efficiency during learning has been previously reported (64). This would prevent the novel social stimulus or any unfamiliar or unrelated stimuli from modifying overall network topology during a state of reconsolidation (63). Whether neural activity involved in reconsolidation of prior information can inhibit a consolidation of newly encountered information was not directly assessed in the present design. We do take note. However, that the applied graph algorithms may have not been sensitive to modality-specific changes in functional networks. Future refinements to stimulus presentations at short time delays prior to scanning may better isolate memory versus non-memory effects on functional networks.

The brain regions shown to have greater node strength and eigenvector centrality in control relative to stimulus exposed rats have been studied in part for their roles in learning and memory. Specifically, these regions may play significant roles in spatial learning, and affective and episodic memories. However, their roles in social learning and recognition are not established. Parietal cortex is known to play important roles in long-term spatial memory functions in rodents and primates (65). The anterior cingulate is involved in establishment of long-term valence associations, such as pain and reward (66, 67). Hippocampally-driven reconsolidation processes may involve communication with retrosplenial cortical circuits for long-term storage (68, 69). Insular cortex plays an important role in memory of aversive experiences (70, 71). It also plays significant roles in reward based learning as does the striatum (72–74). The somatosensory cortex shares input specific synaptic inputs with the hippocampus related to sensory-memory consolidation (75).

Reduced network efficiency and clustering are inversely related to an increase in the average number of intermediate steps separating any pair of nodes in a network (increased characteristic path length). We note that this should not be taken here to imply anatomical steps or axonal connections. Instead, we infer from these network quantifiers variations in the degree of correlated BOLD activity across regions. Regions with highest statistical correlations in BOLD thus represent shortest routes of communication compared with regions with intermediate, low, or weakest pairwise correlations (21, 63). A short (and efficient) path would be characterized by regions linked by the highest or strongest statistical correlations in BOLD. Although an underlying neurophysiological mechanism remains unclear, synchronous activity plays a central role in the temporal entrainment of spike delays to theta rhythms in distributed neuronal populations communicating between cortical regions and hippocampus (76–78). Such spike-timing adjustments to behaviorally-driven oscillations could play a significant role in information transfer and long-term memory and plasticity (63). As indicated above, a change in rhythmic activity may impact band limited power of specific frequencies in cortex and this has the potential of desynchronizing BOLD activity when shifting from a baseline state of rest to a state where a behavioral task was carried out shortly before imaging. At this time, the mechanisms remain unclear. However, there is support in the literature for a reduced global efficiency associated with high rates of learning (64).

We further assessed quantifiers that while not having solid grounding in known neurophysiological mechanisms, may help deepen our understanding of the structure of networks in relation to their efficiency. Thus, we did not observe differences in the degree of difficulty finding edge sequences that form random shortest paths between nodes (search information) nor the number of edges that constitute detours in randomly transited paths between nodes (path transitivity). We did observe that edges along single short ‘hops’ or individual steps between two nodes constituted slightly higher efficient routes for information transfer in control relative to context exposed rats. This is likely associated with a higher number of close local connections (higher clustering coefficient) with short distances (inversely related to high correlation coefficients edge-connected nodes) in controls relative to context-exposed groups. In sum, calculations of shortest path lengths and efficiency may reveal interesting and novel views of the mechanisms of learning and memory, both in terms of organization of edge weights and nodal connectivity patterns involved. Further research into links between efficiency quantifiers and underlying neurophysiological and molecular/cellular correlates are likely to improve the modeling of cognitive mechanisms in animal models.

Aside from the effects of familiar context exposure in reducing node strength, clustering, efficiency, and hub locations, we failed to observe effects of the stimulus on a range of additional network metrics. Contrary to previous results in human subjects undergoing specific cognitive and learning tasks (79, 80), we did not observe changes in global modularity in response to familiar or novel stimuli. Our analysis of node roles also indicated no overall differences in nodal cartographic assignments according to Guimares and Amaral (42) or more recently in functional networks by Meunier and colleagues (43). Consistent with a lack of differences in modular structures, we also observed no differences in assortativity and rich club indices (suggesting no differences in degree distributions). As a way to assess whether changes in node strength and hub reallocations can impact network robustness, we used a targeted node elimination strategy. No differences in network robustness were observed either in response to graded removal of high-to-low strength nodes, or betweenness or eigenvector centrality nodes (we also removed nodes according to their degree values and observed no differences). A random failure model also indicated no differences in overall network robustness (data not shown). A close examination of composite 3D connectome maps in which the nodes were weighted by eigenvector centrality values (Figure 5E) showed stark differences in nodal sizes between control and context-exposed groups. In the latter, node sizes for deep subcortical structures in the hypothalamus appear enlarged. The cortical areas with highest centrality in control rats were the same regions with reduced node strength in context-exposed groups. This suggested that these regions were specifically involved in the reorganization of BOLD correlations induced by the familiar contextual stimulus. The role of hypothalamic structures in the context groups is somewhat surprising as these regions are usually not included as seed regions in functional connectivity studies. Deeper structures are subject to large blood vessel pulsations that are typically well above the resting state frequency power. Undersampled Nyquist frequencies can alias into the measured fMRI signal and drive hypothalamic ‘synchronous’ activity in the presence of suppressed cortical connectivity strength. However, despite this possible non-neuronal mechanism there are studies that have observed a role for hypothalamic nuclei in functional connectivity in models of pain, in human studies of migraine headache, and in studies of obesity where increases in hypothalamic functional connectivity have been reported (81–83).

We reanalyzed subgroups of male rats that were imaged under the same experimental conditions. A subset of these rats were handled and given an intraperitoneal sterile injection 10 minutes before anesthetic induction and preparations for imaging. As a comparison, we also analyzed a group of rats undergoing the exact same injection procedure 24 hours before imaging and a naive group never administered any injection. Results from this analysis indicated that a saline injection before imaging caused a robust increase in connectivity strength, along with other quantifiers that strongly supported a general increase in functional connectivity involving, to a large extent, cortical and thalamic areas. It is interesting to note that the strong salience of this injection stimulus prior to scanning enhanced the ability of a novel stimulus to modify functional connectivity (over any other contextual conditions). One possibility for such a phenomenon might be the release of corticosterone which can act upon memory networks via hippocampal, insular, and basolateral amygdala neurons (84–86) . The importance of this ‘positive control’ study is that it offers some evidence that the changes observed in response to the familiar context stimuli and the novel social stimulus are not solely due to a general or non-specific arousal state. However, an underlying neurobiological mechanism for the observed differences caused by the saline injection also remains to be investigated. One possibility may be the rapid release of catecholamines (87) which may act upon distributed regions of the cortex in a less input selective manner compared with excitatory glutamatergic inputs (21). This can be studied using pre-scan stimulus paradigms involving unexpected versus expected stimuli with or without affective value. One possibility is that ascending cortical noradrenergic inputs from brainstem are activated by the unexpected and arousing injection procedure. These animals did have stable respiratory rates and arterial oxygen saturation (23). Therefore, systemic physiological instability in unlikely to have caused the increases in overall network strength.

We should note several limitations of the present study. First, sedatives and anesthetics exert various effects on functional connectivity and BOLD signal activity based on their underlying molecular targets (88). These include the magnitude of the BOLD signal, the strength of correlations (59), the specificity of connectivity patterns, and these are related in part with the direct actions of isoflurane in blood vessel smooth muscle cells and small arteriolar vessel relaxation as well as the neuronal actions of this anesthetic (89). Use of dexmedetomidine is currently more widely accepted than use of isoflurane but it is no less inert in altering baseline conditions via actions on cerebrovascular parameters and neuronal activity mechanisms. We have conducted awake rat imaging in the past in the context of cue stimulated reward associations (90), but the experience related to awake acclimatization involving repeated restraint sessions and repeated isoflurane inductions would alter the background upon which the contextual and social stimuli were presented. We should, however, note that while the animal is under low sedation and low level anesthesia during scanning, the stimulus is presented with a temporal delay that would allow the generation of neuronal responses and preservation of some degree of network topology prior to setup and scanning. A second limitation is the non-specific simplified stimulus presentation model and its loose association with memory mechanisms. With regards to the association of the current stimulus paradigm with memory, it is important to note that rats were exposed over 3 days to the same test cage and then placed into the same test cage for 10 minutes prior to scanning. This pre-scan experience resulted in robust changes in node strength and other connectome quantifiers relative to unexposed animals. The same outcome was observed in two groups of rats undergoing the same context-exposure regime, those without any additional stimuli and those also presented with a novel unfamiliar social stimulus rat. This supports the possibility that the effect of exposing rats to the pre-scan familiar context was reproducible within the same cohort of rats, both male and females. We should also note that the word ‘memory’ is used here to denote the preservation of a pattern of network topology in stimulus groups that is absent in control unexposed groups. The paradigm used here did not generate a memory ‘response’ by the rats prior or after scanning and this weakens the direct linkage between the network differences between groups and their role in memory structure in the brain. Regarding the experimental paradigm itself, we have previously applied better validated and more system selective experimental paradigms for assessing learning and memory that allowed *a priori* hypothesis testing of the brain regions involved in memory processing (18, 19). We anticipate future studies using similar paradigms here but with the added modification that the memory-retrieving process or the memory-signaling behavior be carried out or measured during a short temporal delay prior to imaging. Finally, approaches to selectively modify the activity within neuronal circuits and drive network differences between control and stimulus groups were not conducted here. Optogenetic and chemogenetic approaches along with direct deep brain electrical stimulation have been applied to study the function of hippocampal circuitry in relation to broader network topology has been studied previously (91). Our objective in future studies will be to integrate variants of these approaches to quantitative assessments of memory networks in control and in aging or neurological disease rodent models.

The present study is an initial step towards understanding the anatomical complexity and dynamic nature of memory in a rat model. The use of ultrahigh field imaging along with connectome analyses provide the tools and allows room for future improvement of the spatial and temporal imaging acquisitions and in the precision and selectivity of network quantifiers to further investigate mechanisms of memory. A major goal of future studies will be to determine the reproducibility of these quantifiers and characterize these as potential biomarkers for basic mechanisms of memory, aging-associated decline in learning and memory, and the impact of neurodegenerative changes on the topology of memory networks.

## Acknowledgments

Supported by National Institute on Drug Abuse grant R03 DA042971, R21 AG065819 and University of Florida College of Medicine Opportunity fund (DRPD-ROF2019) to MF. LMC-P was supported by a McKnight Brain Foundation postdoctoral fellowship and is supported by K25 DA047458 and a young investigator grant from the Brain and Behavior Research Foundation. The contents of this manuscript are solely the responsibility of the authors and do not necessarily represent the official views of the funding agencies. This work was performed in the McKnight Brain Institute at the National High Magnetic Field Laboratory’s AMRIS Facility, which is supported by National Science Foundation Cooperative Agreement No. DMR-1644779 and the State of Florida.

## Notes

### Competing Interest Statement

The authors have declared no competing interest.

### Summary of Updates

In this new revision we edited to the final version being submitted for peer review. We found further minor grammatical errors and edited several citations that were no longer needed.

